# CDK-Mediated Phosphorylation of FANCD2 Promotes Mitotic Fidelity

**DOI:** 10.1101/2020.09.29.318055

**Authors:** Juan A. Cantres-Velez, Justin L. Blaize, David A. Vierra, Rebecca A. Boisvert, Jada M. Garzon, Benjamin Piraino, Winnie Tan, Andrew J. Deans, Niall G. Howlett

## Abstract

Fanconi anemia (FA) is a rare genetic disease characterized by increased risk for bone marrow failure and cancer. The FA proteins function together to repair damaged DNA. A central step in the activation of the FA pathway is the monoubiquitination of the FANCD2 and FANCI proteins under conditions of cellular stress and during S-phase of the cell cycle. The regulatory mechanisms governing S-phase monoubiquitination, in particular, are poorly understood. In this study, we have identified a CDK regulatory phospho-site (S592) proximal to the site of FANCD2 monoubiquitination. FANCD2 S592 phosphorylation was detected by LC-MS/MS and by immunoblotting with a S592 phospho-specific antibody. Mutation of S592 leads to abrogated monoubiquitination of FANCD2 during S-phase. Furthermore, FA-D2 (*FANCD2^-/-^*) patient cells expressing S592 mutants display reduced proliferation under conditions of replication stress and increased mitotic aberrations, including micronuclei and multinucleated cells. Our findings describe a novel cell cycle-specific regulatory mechanism for the FANCD2 protein that promotes mitotic fidelity.

**Author Summary:** Fanconi anemia (FA) is a rare genetic disease characterized by high risk for bone marrow failure and cancer. FA has strong genetic and biochemical links to hereditary breast and ovarian cancer. The FA proteins function to repair DNA damage and to maintain genome stability. The FANCD2 protein functions at a critical stage of the FA pathway and its posttranslational modification is defective in >90% of FA patients. However, the function, and regulation of FANCD2, particularly under unperturbed cellular conditions, remains remarkably poorly characterized. In this study, we describe a novel mechanism of regulation of the FANCD2 protein during S-phase of the cell cycle. CDK-mediated phosphorylation of FANCD2 on S592 promotes the ubiquitination of FANCD2 during S-phase. Disruption of this phospho-regulatory mechanism results in compromised mitotic fidelity and an increase in mitotic chromosome instability.

## Introduction

Protection of the integrity of our genome depends on the concerted activities of several DNA repair pathways that ensure the timely and accurate repair of damaged DNA. These DNA repair pathways need to be tightly coordinated and regulated upon cellular exposure to exogenous DNA damaging agents, as well as during the cell cycle. Somatic disruption of the DNA damage response leads to mutation, genome instability and cancer. In addition, germline mutations in DNA repair genes are associated with hereditary diseases characterized by increased cancer risk and other clinical manifestations. Fanconi anemia (FA) is a rare genetic disease characterized by congenital abnormalities, increased risk for bone marrow failure and cancer, and accelerated aging [1]. FA is caused by germline mutations in any one of 23 genes. The FA proteins function together in a pathway to repair damaged DNA and to maintain genome stability. A major role for the FA pathway in the repair of DNA interstrand crosslinks (ICLs) has been described [2].

The main activating step of the FA pathway is the monoubiquitination of the FANCD2-FANCI heterodimer (ID2), which is catalyzed by the FA-core complex, comprising FANCA, FANCB, FANCC, FANCE, FANCF, FANCG, FANCL, UBE2T/FANCT, and the FA associated proteins. Monoubiquitination occurs following exposure to DNA damaging agents and during S-phase of the cell cycle [3, 4]. Recently, it was discovered that the monoubiquitination of FANCD2 and FANCI leads to the formation of a closed ID2 clamp that encircles DNA [5, 6]. Moreover, ubiquitinated ID2 (ID2-Ub) assembles into nucleoprotein filament arrays on double-stranded DNA [7]. The monoubiquitination of FANCD2 is necessary for efficient interstrand crosslink repair, the maintenance of common fragile site stability and faithful chromosome segregation, events that are all crucial for genomic stability [8, 9].

A DNA damage-independent role for the FA pathway during the cell cycle has been implicated in several studies. For example, FANCD2 promotes replication fork protection during S-phase and ensures the timely and faithful replication of common chromosome fragile sites (CFSs) [8, 10, 11]. FANCD2 has also been shown to be involved in mitotic DNA synthesis (MiDAS) during prophase and is present on the terminals of anaphase ultrafine bridges [12, 13]. During the cell cycle FANCD2 monoubiquitination is maximal during S-phase and minimal during M-phase [4]. FANCD2 and FANCI are phosphorylated by the ATR and ATM kinases following exposure to DNA damaging agents, promoting their monoubiquitination [14–17]. However, in general, the function and regulation of FANCD2 and FANCI during the cell cycle remains poorly understood.

Cyclin-dependent kinases (CDKs) play a major role in regulating cell cycle progression, with CDK-mediated hyperphosphorylation of the retinoblastoma protein (pRb) being the primary mechanism of cell cycle regulation. CDKs are also known to regulate DNA repair during the cell cycle. Several protein components of the homologous recombination repair (HR), translesion DNA synthesis (TLS), non-homologous end joining (NHEJ), and telomere maintenance pathways are CDK substrates [18]. For example, the BRCA2 protein is phosphorylated by CDK1 as cell progress towards mitosis. CDK-mediated phosphorylation of BRCA2 blocks interaction with RAD51 and thereby restricts HR DNA repair during M-phase [19]. Conversely, CDK1/2 phosphorylate the EXO1 nuclease to promote DNA strand resection and HR repair during S-phase [20].

In this study we have examined the regulation of the FANCD2 protein by phosphorylation, specifically during the cell cycle. We show that FANCD2 is phosphorylated on S592, a putative CDK site, during S-phase but not during M-phase and is phosphorylated by CDK2 *in vitro*. Mutation of S592 as well as CDK inhibition disrupts S-phase FANCD2 monoubiquitination. Furthermore, we demonstrate that FA-D2 (*FANCD2^-/-^*) patient cells stably expressing FANCD2 mutated at S592 display reduced growth under conditions of replication stress, an altered cell cycle profile, and increased genomic instability manifested as increased micronuclei, nucleoplasmic bridges, and aneuploidy. Our results uncover important mechanistic insight into the regulation of a key DNA repair/genome maintenance pathway under unperturbed cellular conditions.

## Results

### FANCD2 is phosphorylated under unperturbed conditions and during S-phase of the cell cycle

To study the phosphorylation of the FANCD2 protein in the absence and presence of DNA damaging agents, we performed a lambda-phosphatase assay with several cell lines following incubation in the absence or presence of the DNA crosslinking agent mitomycin C (MMC). Notably, we observed a large increase in FANCD2 mobility following incubation of lysates with lambda-phosphatase even in the absence of MMC in all cell lines examined (Fig. 1A). These results suggest that FANCD2 is subject to extensive phosphorylation even in the absence of an exogenous DNA damaging agent. A similar change in protein mobility was not observed for FANCI (Fig. 1A). To determine if FANCD2 is subject to phosphorylation during the cell cycle, we performed a double-thymidine block experiment causing an early S-phase arrest and analyzed phosphorylation at regular time points following release. We observed maximal phosphorylation of FANCD2 during S-phase of the cell cycle, with much less phosphorylation observed as cells progressed through G2/M-and G1-phases of the cell cycle (Fig. 1B). Again, we observed no appreciable change in FANCI mobility under the same conditions, suggesting that FANCI is not subject to the same levels of phosphorylation as FANCD2 during the cell cycle (Fig. 1B). Similar findings were observed with U2OS cells (Fig. S1A and B). We also synchronized cells in M-phase using nocodazole and, upon release, again observed maximal levels of FANCD2 phosphorylation during S-phase (∼15 h after release) (Fig. S1C and D). The FANCA protein is a central component of the FA core complex, a multi-subunit ubiquitin ligase that catalyzes the monoubiquitination of FANCD2 [3, 21]. To determine if FANCD2 phosphorylation was dependent on its monoubiquitination, we performed a lambda-phosphatase assay with asynchronous and early S-phase synchronized FA-A (*FANCA^-/-^*) and FANCA-complemented FA-A cells. S-phase FANCD2 phosphorylation was observed in the absence of FANCA, albeit to a slightly lesser extent than in cells FANCA-complemented FA-A cells (Fig. 1C). These results suggest that S-phase phosphorylation is coupled to, but not dependent upon, the monoubiquitination of FANCD2.

**Figure 1.**
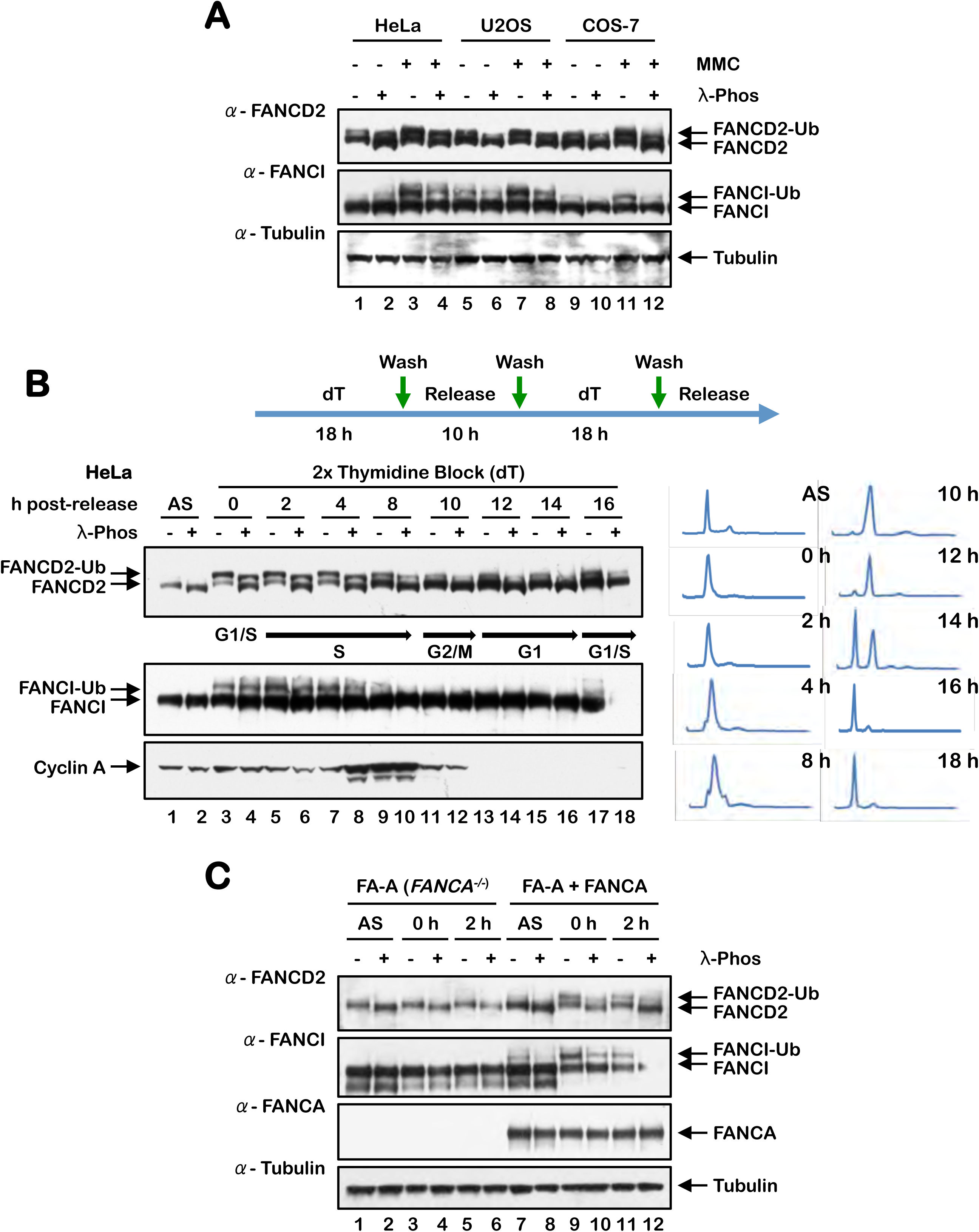
FANCD2 is phosphorylated under non-stressed conditions and during S-phase of the cell cycle. (A) HeLa, U2OS and COS-7 cells were incubated with or without 200 nM MMC for 24 h. Cells were harvested and lysed in lambda phosphatase lysis buffer, incubated in the absence or presence of lambda phosphatase, and the indicated proteins were analyzed by immunoblotting. (B) HeLa cells were synchronized at the G1/S boundary by a double-thymidine block and released into thymidine-free media. Cells were lysed in lambda phosphatase buffer, incubated in the presence or absence of lambda phosphatase, and lysates analyzed by immunoblotting. For cell cycle stage analysis, cells were fixed, stained with propidium iodide and analyzed by flow cytometry. (C) FA-A (*FANCA^-/-^*) and FANCA-complemented FA-A cells were synchronized at the G1/S boundary by a double-thymidine block and released into thymidine-free media. Cells were harvested, lysed in lambda phosphatase buffer, incubated in the presence or absence of lambda phosphatase, and analyzed by immunoblotting.

### FANCD2 is a CDK substrate

Phosphorylation of FANCD2 during the cell cycle suggested a role for cyclin-dependent kinases (CDKs) in the regulation of FANCD2. CDKs phosphorylate serine or threonine residues in the consensus sequence [S/T*]PX[K/R], and several DNA repair proteins are known CDK substrates, including BRCA2, FANCJ, and CtIP [19, 22, 23]. Both FANCD2 and FANCI have several putative CDK phosphorylation sites with varying degrees of conservation (Fig. S2A and B). To begin to assess the role of CDKs in the regulation of FANCD2, we performed a double-thymidine block in the absence and presence of the CDK inhibitors purvalanol A and SNS-032. At the concentrations tested, treatment with both inhibitors resulted in a significant reduction in the levels of FANCD2 and FANCI monoubiquitination observed after double-thymidine arrest and a modest reduction in FANCD2 phosphorylation (Fig. 2A). Reduced FANCD2 phosphorylation was also observed upon incubation with other CDK inhibitors and upon short-term exposure to purvalanol A (Fig. S2C and D). To determine if FANCD2 is a CDK substrate, we immunoprecipitated FANCD2 from FA-D2 (*FANCD2^-/-^*) patient cells stably expressing V5-tagged LacZ or FANCD2, and probed immune complexes with a pan anti-pS/T-CDK antibody. An immune reactive band was detected in immune complexes from cells expressing FANCD2 and not LacZ (Fig. 2B), suggesting that FANCD2 is a CDK substrate. We also performed an *in vitro* CDK kinase assay with CDK2-Cyclin A and full length FANCD2 purified from High Five insect cells [24]. CDK-mediated FANCD2 phosphorylation was already evident upon purification from insect cells (Fig. 2C). Following dephosphorylation with lambda phosphatase, we observed a concentration-dependent increase in FANCD2 phosphorylation upon incubation with CDK2-Cyclin A. These results indicate that FANCD2 is a CDK substrate and suggest that FANCD2 monoubiquitination may be regulated or coupled with CDK phosphorylation.

**Figure 2.**
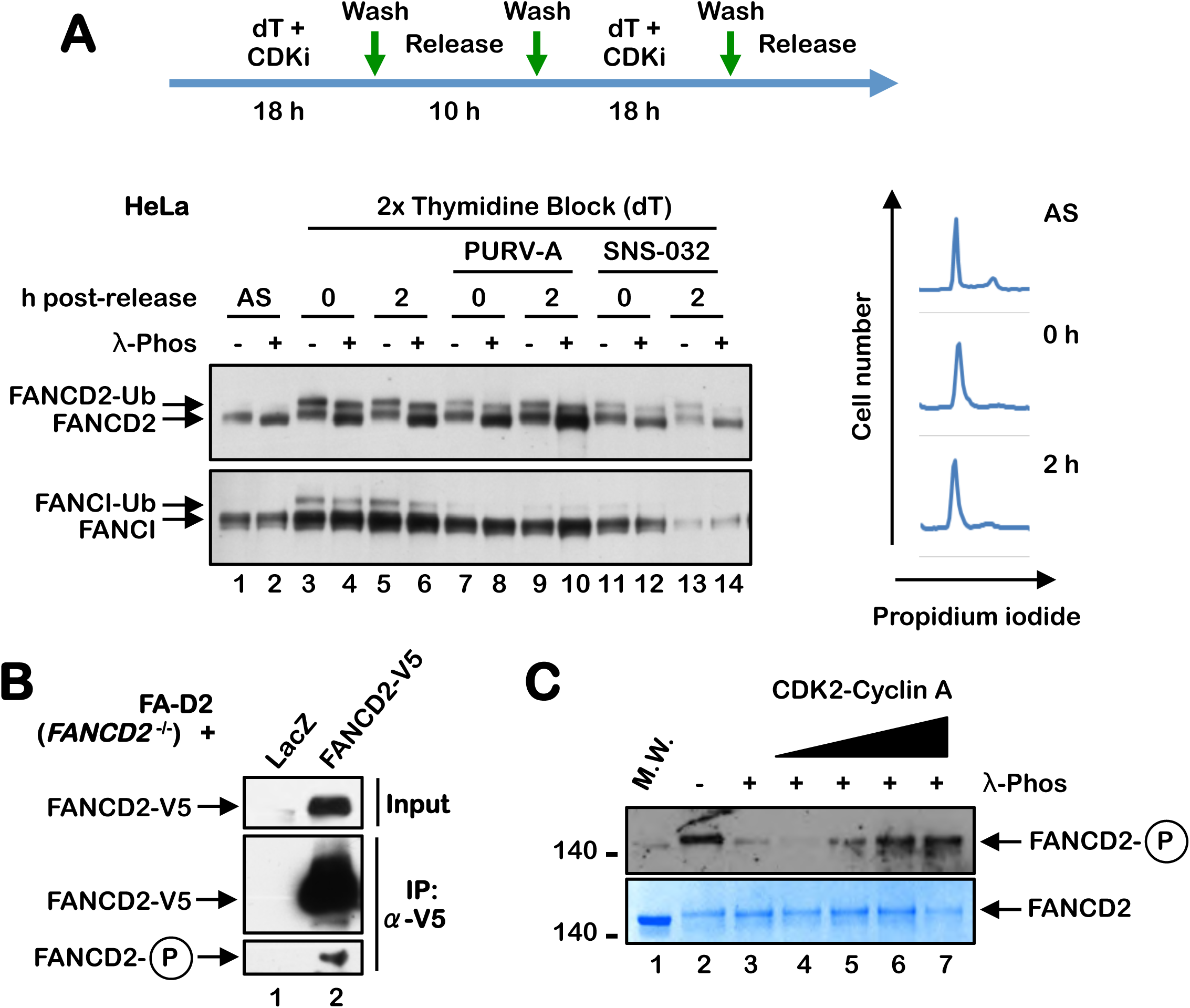
FANCD2 is a CDK substrate. (A) HeLa cells were synchronized at the G1/S boundary by a double-thymidine block performed in the absence or presence of the CDK inhibitors Purvalanol A (PURV-A) or SNS-032. Cells were lysed in lambda phosphatase buffer, incubated in the absence or presence of lambda phosphatase, and analyzed by immunoblotting. Cells were also fixed, stained with propidium iodide and analyzed by flow cytometry to determine cell cycle stage. (B) FA-D2 (*FANCD2^-/-^*) cells stably expressing LacZ-V5 or FANCD2-V5 were immunoprecipitated with anti-V5 agarose and immune complexes were immunoblotted with anti-FANCD2 and anti-pS-CDK antibodies. (C) Purified FANCD2 was incubated in the absence (-) or presence (+) of lambda phosphatase followed by incubation with increasing concentrations of CDK2-Cyclin A and ATP at 30°C for 30 min. Samples were immunoblotted with a pan anti-pS-CDK antibody or stained with SimplyBlue SafeStain.

### FANCD2 is phosphorylated on S592 during S-phase of the cell cycle

To map the *in vivo* sites of FANCD2 phosphorylation, we immunoprecipitated FANCD2 from asynchronous U2OS cells stably expressing 3xFLAG-FANCD2 under stringent conditions, and performed phosphoproteomic analysis using LC-MS/MS (Fig. 3A and B). Under these conditions, we observed the phosphorylation of multiple sites including the previously detected ATM/ATR phosphorylation sites S1401 and S1404 (Table 1) [17]. We also detected phosphorylation of the putative CDK site S592 (Table 1). To analyze the phosphorylation of FANCD2 during the cell cycle, we synchronized HeLa cells in M-phase using nocodazole, immunoprecipitated FANCD2 immediately upon release and at 15 h post-release and again performed phosphoproteomic analysis using LC-MS/MS (Fig. 3C-F). Under these conditions, we detected phosphorylation of FANCD2 on S592 in S-phase and not during M-phase. To study FANCD2 S592 phosphorylation more closely, we generated a S592 phospho-specific antibody. We arrested cells in M-phase using nocodazole and analyzed FANCD2 S592 phosphorylation upon release from nocodazole block. A FANCD2 pS592 immunoreactive band was observed at 12 h (S-phase) following release (Fig. 3G), consistent with the results of our LC-MS/MS phosphoproteomic analysis (Fig. 3F, Table 1). Taken together our results suggest that the phosphorylation of FANCD2 S592 might play an important regulatory function during S-phase (Fig. 3F).

**Figure 3.**
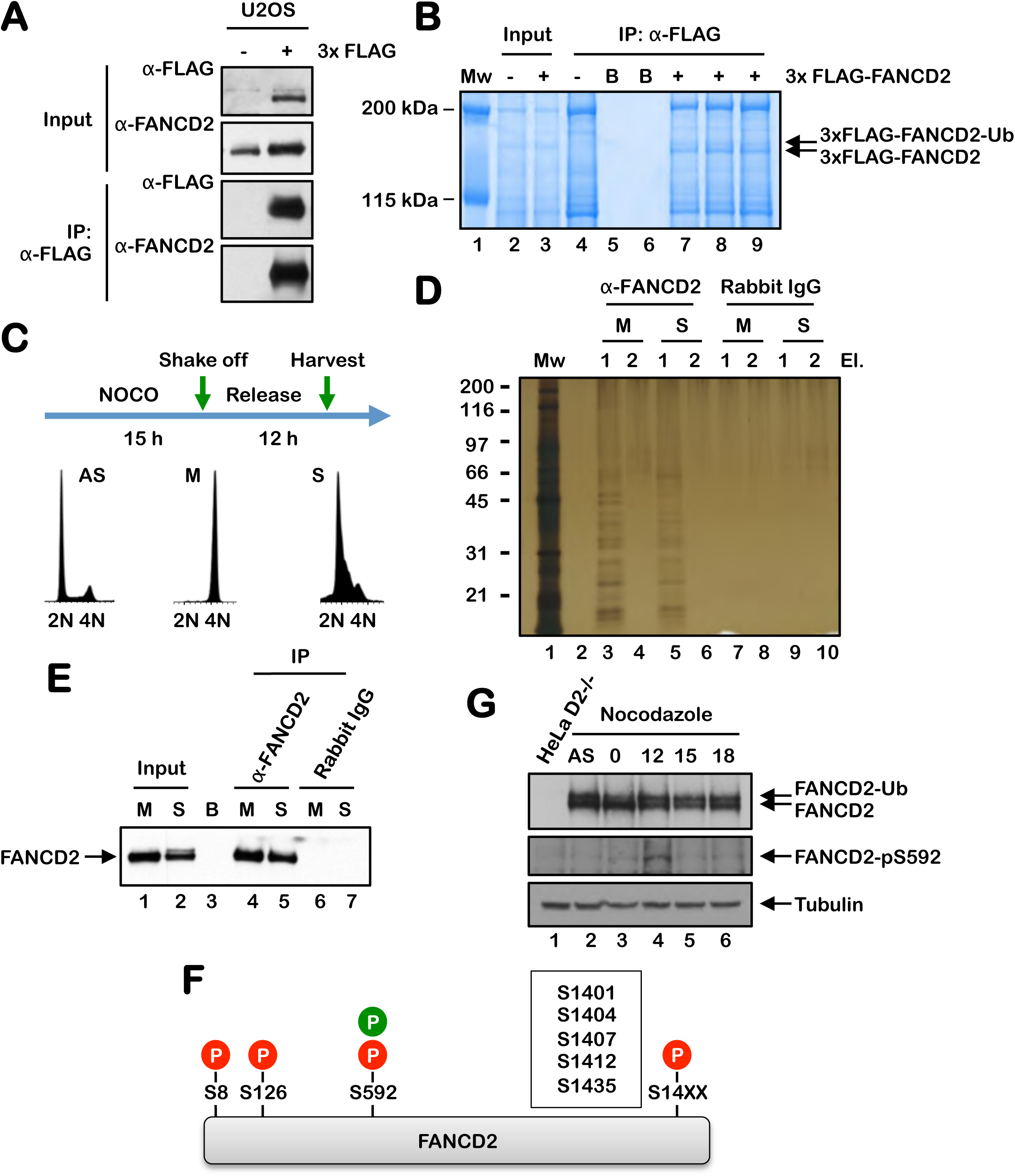
FANCD2 is phosphorylated on S592 during S-phase. (A-B) FANCD2 was immunoprecipitated from U2OS cells stably expressing 3xFLAG-FANCD2 under stringent conditions using anti-FLAG agarose. Immune complexes were analyzed by immunoblotting using anti-FLAG and anti-FANCD2 antibodies (A), and by staining with SimplyBlue SafeStain (B). Immunoprecipitated FANCD2 bands were combined and subjected to phosphoproteomic analysis using LC-MS/MS. (C) HeLa cells were synchronized in M-phase by nocodazole block. Mitotic cells were harvested or released into nocodazole-free medium for 12 h (S-phase) prior to harvesting. (D) FANCD2 was immunoprecipated from M-phase and S-phase synchronized cells using an anti-FANCD2 antibody, and visualized using silver staining (D) or by immunoblotting (E). (F) A schematic of the FANCD2 protein indicating phosphorylation sites identified by LC-MS/MS. (G) HeLa cells were synchronized in M-phase by nocodazole block. Samples were harvested and analyzed by immunoblotting using an anti-FANCD2 antibody and an anti-FANCD2 pS592 phospho-specific antibody.

### Mutation of S592 disrupts S-phase FANCD2 monoubiquitination

FANCD2 S592 is predicted to localize to a flexible loop proximal to K561, the site of monoubiquitination, suggesting a potential regulatory function for S592 phosphorylation (Fig. 4A). To begin to characterize the role of S592 phosphorylation, we generated phospho-dead (S592A) and phospho-mimetic (S592D) variants and stably expressed these in FA-D2 (*FANCD2^-/-^*) patient-derived cells (Fig. 4B). Mutation of S592 did not adversely impact the stability or overall structure of FANCD2 as evidenced by equal protein expression levels and intact monoubiquitination following MMC exposure (Fig. S3A). Previous studies have shown that FANCD2 undergoes monoubiquitination as cells progress through S-phase of the cell cycle [4]. To analyze the effects of S592 mutation on S-phase FANCD2 monoubiquitination, cells were subject to a double-thymidine block, and FANCD2 monoubiquitination was analyzed upon release. Consistent with previous studies, we observed an increased in monoubiquitination of wild-type FANCD2 as cells progressed through S-phase (Fig. 4C). In contrast, S-phase monoubiquitination was markedly attenuated for both FANCD2-S592A and FANCD2-S592D (Fig. 4C). We also analyzed levels of the mitotic marker H3 pS10 in these cells. Compared to cells expressing wild-type FANCD2, we observed persistent levels of H3 pS10 in FA-D2 cells expressing empty vector and the FANCD2-S592A mutant (Fig. 4C). We observed a similar phenotype of elevated H3 pS10 in HeLa *FANCD2^-/-^* cells generated by CRISPR-Cas9 gene editing (Fig. S3B-D) [25]. In contrast to cells expressing empty vector and the FANCD2-S592A mutant, we observed a more rapid disappearance of H3 pS10 in cells expressing FANCD2-S592D (Fig. 4C). In addition, FACS analysis of FA-D2 patient cells expressing empty vector and the S592 variants upon release from double-thymidine block indicated a higher percentage of cells in G2/M at earlier time points, compared to cells expressing FANCD2-WT (Fig. S3E). Taken together, these results demonstrate that mutation of FANCD2 S592 disrupts S-phase monoubiquitination and leads to altered G2-M cell cycle progression.

**Figure 4.**
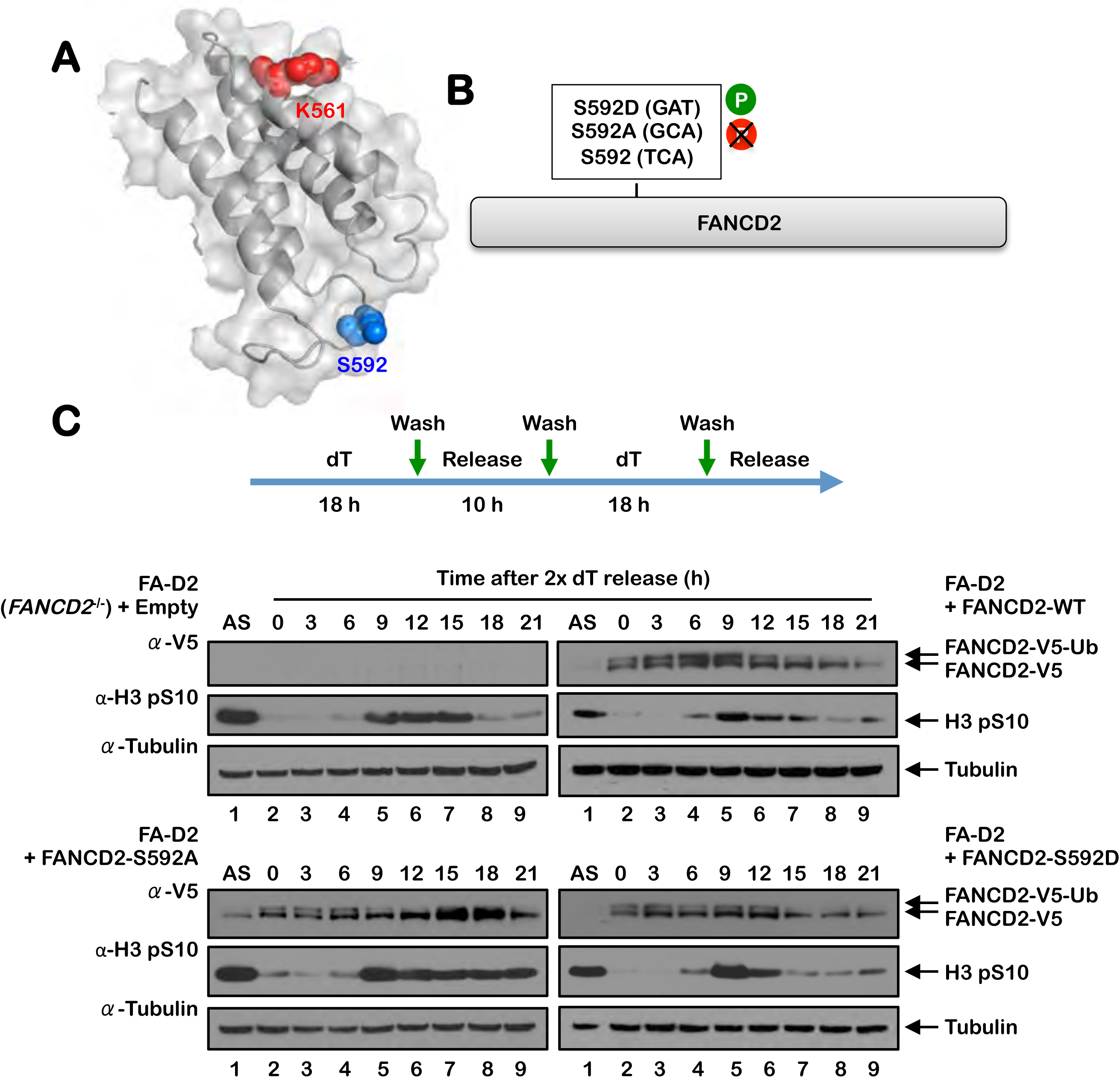
Mutation of S592 disrupts FANCD2 monoubiquitination during S-phase and following exposure to DNA damaging agents. (A) Shown is a partial model of the human FANCD2 protein, modeled on the mouse Fancd2 structure (PDB #3S4W), illustrating the site of monoubiquitination K561 in red and S592 in blue. (B) Site-directed mutagenesis was used to generate phospho-dead (S592A) and phospho-mimetic (S592D) FANCD2 S592 variants. (C) FA-D2 cells stably expressing empty vector or V5-tagged FANCD2-WT, FANCD2-S592A, or FANCD2-S592D were synchronized at the G1/S boundary by double-thymidine block. Cells were released into thymidine-free medium and whole cell lysates were analyzed by immunoblotting at the indicated times post-release.

### Mutation of FANCD2 S592 leads to decreased proliferative capacity and increased mitotic defects

To assess the functional impacts of mutation of FANCD2 S592, we monitored the proliferation of FA-D2 patient cells stably expressing FANCD2-WT and the S592 variants in the presence of low concentrations of DNA damaging agents for prolonged periods using the xCELLigence real time cell analysis system. The xCELLigence system enables the analysis of cellular phenotypic changes by continuously monitoring electrical impedance. Impedance measurements are displayed as the cell index, which provides quantitative information about the biological status of the cells, including cell number, cell viability, and cell morphology. Compared to cells expressing FANCD2-WT, cells expressing empty vector and the S592 variants exhibited reduced proliferation when cultured in the presence of low concentrations of the DNA polymerase inhibitor aphidicolin (APH) (Fig. 5A and S5A). In contrast, mutation of FANCD2 S592 had no discernible impact on growth in the presence of low concentrations of mitomycin C (MMC) (Fig. 5A and S5A). We also analyzed growth in the presence of the CDK1 inhibitor RO3306. CDK1 inhibition resulted in reduced cell proliferation in cells expressing FANCD2-WT and the S592A variant compared to cells expressing empty vector or the S592D variant (Fig. S5B and C). Next, we examined the effects of mutation of S592 on levels of mitotic aberrations. Compared to FA-D2 cells expressing wild-type FANCD2, we observed increased levels of micronuclei, nucleoplasmic bridges, and bi- and multi-nucleated cells in FA-D2 cells expressing the S592 variants (Fig. 5B). Elevated levels of mitotic aberrations were also observed in FA-D2 cells expressing FANCD2-K561R, which cannot be monoubiquitinated (Fig. 5B). Taken together, these results indicate that S592 phosphorylation is functionally necessary to ensure mitotic fidelity and chromosome stability under non-stressed conditions.

**Figure 5.**
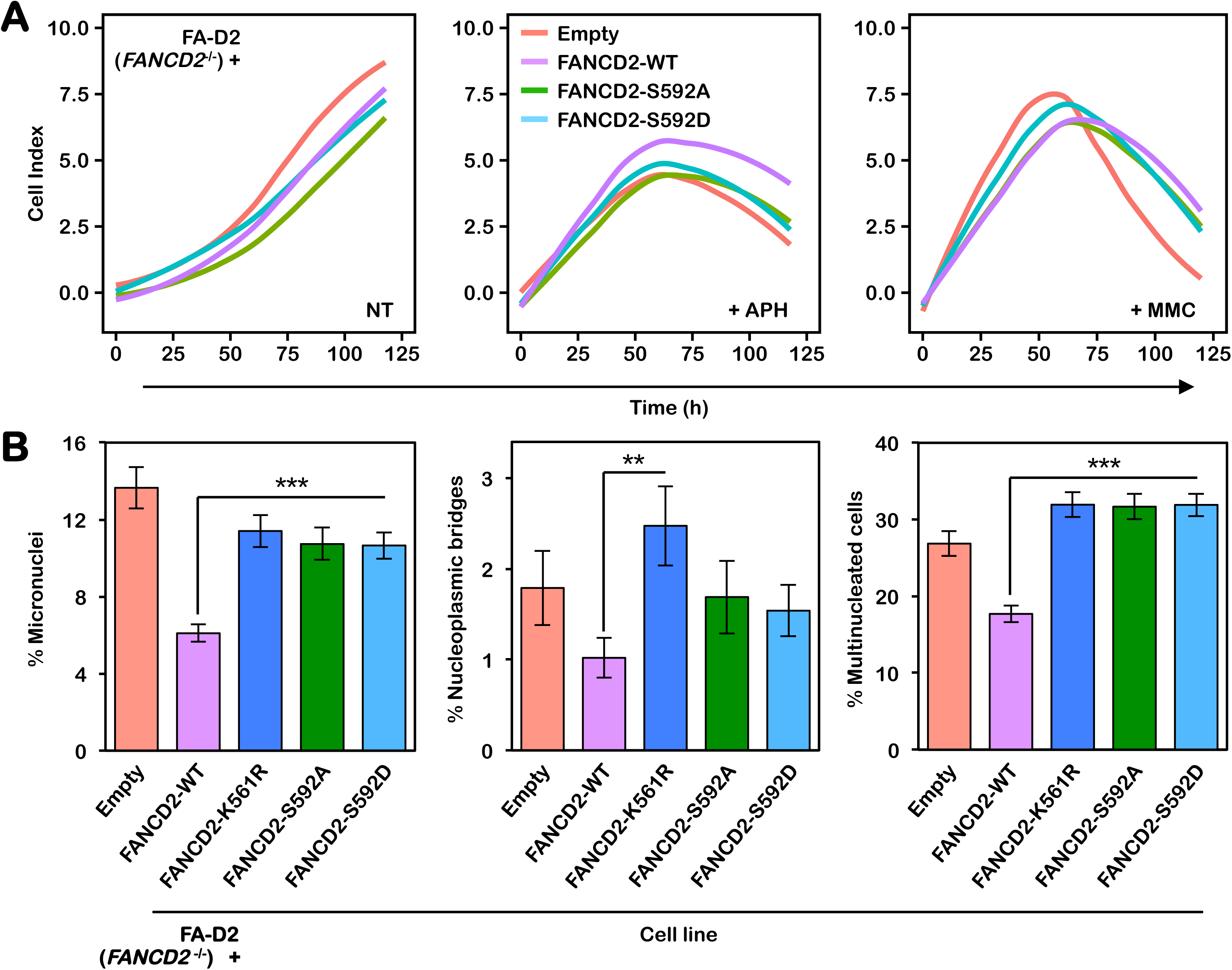
Mutation of FANCD2 S592 leads to decreased proliferation under conditions of replicative stress and increased mitotic defects. (A) FA-D2 cells stably expressing empty vector or V5-tagged FANCD2-WT, FANCD2-S592A, or FANCD2-S592D were incubated in the absence (NT) or presence of 0.4 μM aphidicolin (+APH) or 20 nM mitomycin C (+MMC). Cellular proliferation was monitored by measuring electrical impedance every 15 minutes over a 120 h period using the xCELLigence real time cell analysis system. (B) FA-D2 cells stably expressing empty vector or V5-tagged FANCD2-WT, FANCD2-K561R, FANCD2-S592A, or FANCD2-S592D were incubated in the absence of any DNA damaging agents and mitotic aberrations including micronuclei, nucleoplasmic bridges, and multinucleated (>2) cells were scored. At least 400 cells per group were scored and the combined results from two independent experiments are shown. **, *P* < 0.05; ***, *P* < 0.001.

## Discussion

In this study, we have investigated the posttranslational regulation of the central FA pathway protein FANCD2 largely under unperturbed conditions. The majority of studies to date have focused on the posttranslational regulation of FANCD2 and FANCI following exposure to DNA damaging agents [15, 16, 26]. Here, we have observed that FANCD2 undergoes extensive phosphorylation during the cell cycle and, in particular, during S-phase of the cell cycle. Notably, in contrast, FANCI - a FANCD2 paralog - does not appear to be subject to the same degree of phosphorylation during the cell cycle. We have also determined that FANCD2 is phosphorylated on S592 by CDK2-Cyclin A during S-phase using several approaches, including phosphoproteomic analysis of FANCD2 immune complexes from synchronized cell populations and immunoblotting with a S592 phospho-specific antibody. Previous studies have shown that FANCD2 is monoubiquitinated as cells traverse S-phase, and this has been shown to contribute to the protection of stalled replication forks from degradation [4, 11]. Here we show that mutation of S592, or CDK inhibition, abrogates S-phase monoubiquitination, strongly suggesting that S592 phosphorylation primes FANCD2 for ubiquitination during S-phase. Recent studies have established that the FANCL RING E3 ubiquitin ligase allosterically alters the active site of the UBE2T E2 ubiquitin-conjugating enzyme to promote site-specific monoubiquitination of FANCD2 [27]. Specifically, FANCL binding to UBE2T exposes a basic triad of the UBE2T active site, promoting favorable interactions with a conserved acidic patch proximal to K561, the site of ubiquitination [27]. We speculate that S592 phosphorylation may augment interaction with the basic active site of UBE2T. Alternatively, S592 phosphorylation may inhibit FANCD2 de-ubiquitination by USP1, in a manner similar to that previously reported for the FANCI S/TQ cluster [15].

Our studies further emphasize the critical nature of coordinated posttranslational modification of FANCD2. Several studies have previously established intricate dependent and independent relationships between FANCD2 monoubiquitination and phosphorylation. For example, the ATM kinase phosphorylates FANCD2 on several S/TQ motifs following exposure to ionizing radiation, e.g. S222, S1401, S1404, and S1418 [17] - *phosphorylation of S1401 and S1404 were also detected under unperturbed conditions in this study*. While phosphorylation of S222 promotes the establishment of the IR-inducible S-phase checkpoint, S222 phosphorylation and K561 monoubiquitination appear to function as independent events [17]. In contrast, phosphorylation of FANCD2 on T691 and S717 by the ATR kinase is required for efficient FANCD2 monoubiquitination [16, 28]. CHK1-mediated phosphorylation of FANCD2 S331 also promotes DNA damage-inducible monoubiquitination and its interaction with FANCD1/BRCA2 [29]. Conversely, CK2-mediated phosphorylation of FANCD2 inhibits its monoubiquitination and ICL recruitment [30]. Our results suggest that CDK-mediated phosphorylation of S592 primes FANCD2 for monoubiquitination during S-phase in particular. It remains to be determined if other posttranslational modifications, including phosphorylation at other sites, e.g. S1401 and S1404 detected in this study, also contribute to the regulation of FANCD2 during the cell cycle.

Our work aligns with several previous studies describing the coordination of the DNA damage response with cell cycle progression. For example, BRCA2 carboxy-terminal phosphorylation by CDK2 precludes the binding of RAD51, thereby inactivating HR prior to the onset of mitosis [19]. In contrast, CDK1/2 phosphorylation of EXO1 endonuclease positively regulates DNA strand resection and HR repair during S-phase [20]. Similarly, CDK-mediated phosphorylation of CtIP allosterically stimulates its phosphorylation by ATM, resulting in the promotion of strand resection and HR during S-phase [31]. While a role for FANCD2 in HR has been established [32], it remains to be determined if CDK-mediated FANCD2 S592 phosphorylation promotes HR or other related functions. For example, FANCD2 has been shown to have a DNA repair-independent role in protecting stalled replication forks from MRE11-mediated degradation [11]. We have also previously established an important role for FANCD2 in the maintenance of CFS stability: In the absence of FANCD2, CFS replication forks stall at a greater frequency, a likely consequence of persistent toxic R-loop structures, leading to visible gaps and breaks on metaphase chromosomes [8, 10, 33, 34]. Under conditions of replication stress, CFSs can remain unreplicated until mitosis where replication can be completed through a process referred to as mitotic DNA synthesis (MiDAS), a POLD3- and RAD52-dependent process that shares features with break-induced DNA replication (BIR) [35, 36]. FANCD2 has been shown to play a role in MiDAS [12, 13]. Defects in - or an overreliance upon - this process are likely to lead to chromosome missegregation and mitotic defects, as we have observed upon mutation of S592. Increased mitotic aberrations have previously been observed in primary murine FA pathway-deficient hematopoietic stem cells and FA-patient derived bone marrow stromal cells [9]. Taken together, our results support a model whereby CDK-mediated S592 phosphorylation promotes efficient S-phase monoubiquitination, ensuring the efficient and timely completion of CFS replication and mitotic chromosome stability. A greater understanding of the function and regulation of the FA pathway under physiologically relevant non-stressed conditions will provide greater insight into the pathophysiology of this disease and ultimately lead to improved therapeutic options for FA patients.

## Materials and Methods

### Cell culture and generation of mutant cell lines

HeLa cervical carcinoma and U2OS osteosarcoma cells were grown in Dulbecco’s modified Eagle’s medium (DMEM) supplemented with 10% v/v FBS, L-glutamine and penicillin/streptomycin. HeLa FANCD2^-/-^ cells generated by CRISPR-Cas9 were generously provided by Martin Cohn at the University of Oxford [25]. 293FT viral producer cells (Invitrogen) were cultured in DMEM containing 12% v/v FBS, 0.1 mM non-essential amino acids (NEAA), 1% v/v L-glutamine, 1 mM sodium pyruvate and 1% v/v penicillin/streptomycin. PD20 FA-D2 (FANCD2^hy/-^) patient cells were purchased from Coriell Cell Repositories (Catalog ID GM16633). These cells harbor a maternally inherited A-G change at nucleotide 376 that leads to the production of a severely truncated protein, and a paternally inherited missense hypomorphic (^hy^) mutation leading to a R1236H change. PD20 FA-D2 cells were stably infected with pLenti6.2/V5-DEST (Invitrogen) harboring wild-type or mutant FANCD2 cDNAs. Stably infected cells were grown in DMEM complete medium supplemented with 2 μg/ml blasticidin.

### Immunoprecipitation

FA-D2 cells stably expressing LacZ or V5-tagged FANCD2, U2OS, and U2OS 3xFLAG FANCD2 cells were lysed in Triton-X100 lysis buffer (50 mM Tris.HCl pH 7.5, 1% v/v Triton X-100, 200 mM NaCl, 5 mM EDTA, 2 mM Na3VO4, Protease inhibitors (Roche), and 40 mM β-glycerophosphate) on ice for 15 min followed by sonication for 10 s at 10% amplitude using a Fisher Scientific Model 500 Ultrasonic Dismembrator. Anti-V5 or anti-FLAG-agarose were washed and blocked with NETN100 buffer (20 mM Tris-HCl pH 7.5, 0.1% NP-40, 100 mM NaCl, 1 mM EDTA, 1 mM Na3VO4, 1 mM NaF, protease inhibitors (Roche)) plus 1% BSA and the final wash and resuspension was done in Triton-X100 lysis buffer. Lysates were incubated with agarose beads at 4°C for 2 h with nutating. Agarose beads were then washed in Triton-X100 lysis buffer and boiled in 1× NuPAGE buffer (Invitrogen) and analyzed for the presence of proteins by SDS-PAGE and immunoblotting or stained using Colloidal Blue Staining Kit (Invitrogen) for mass spectrometry.

### Cell cycle FANCD2 immunoprecipitation

S-phase and M-phase synchronized populations of HeLa cells were lysed in lysis buffer (20 mM HEPES, pH 7.9, 1.5 mM MgCl2, 400 mM KCl, 0.2 mM EDTA, 20% glycerol, 0.1 mM DTT, 0.1% NP-40, 1 mM PMSF and complete protease inhibitors). 100 units of benzonase was added per 100 μL of lysis buffer. Protein G magnetic beads (Dynabeads, Novex) were crosslinked with anti-FANCD2 (NB100-182, Novus Biologicals) antibody. An equal volume of no salt buffer (20 mM HEPES, pH7.9, 1.5 mM MgCl2, 0.2 mM KCl, 0.2 mM EDTA, 0.1mM DTT, 0.1% NP-40, plus complete protease inhibitors) was added to 2 mg of whole cell lysate. Samples were resuspended in anti-FANCD2-bound beads rotating at 4°C for 4 h. Beads were washed in wash buffer (20 mM HEPES, pH 7.9, 1.5 mM MgCl2, 0.2 mM KCl, 300 mM KCl, 10% glycerol, 0.2 mM EDTA, 0.1 mM DTT, 0.1% NP-40, plus complete protease inhibitor). The samples were eluted in urea elution buffer (8 M urea, 1 mM Na3VO4, 2.5 mM Na₂H₂P₂O, 1 mM β- glycerophosphate and 25 mMHEPES, pH 8.0). Silver staining was performed to visualize the IP and FANCD2 pulldown was confirmed by western blot. Samples were sent for LC-MS/MS analysis at the COBRE Center for Cancer Research Development Proteomics Core at Rhode Island Hospital.

### Lambda phosphatase assay

Cells were harvested and pellets were split into two, lysed in either lambda phosphatase lysis buffer (50 mM Tris-HCl pH 7.5, 150 mM NaCl, 0.1 mM EGTA, 1 mM dithiothreitol, 2 mM MnCl2, 0.01% Brij35, 0.5% NP-40, and protease inhibitor) or lambda phosphatase buffer with the addition of phosphatase inhibitors, 2 mM Na3VO4 and 5 mM NaF for 15 min at 4°C followed by sonication for 10 s at 10% amplitude using a Fisher Scientific Model 500 Ultrasonic Dismembrator. Lysates were incubated with or without 30 U of lambda phosphatase 1 h at 30°C.

### Immunoblotting

For immunoblotting analysis, cell pellets were washed in PBS and lysed in 2% w/v SDS, 50 mM Tris-HCl, 10 mM EDTA followed by sonication for 10 s at 10% amplitude. Proteins were resolved on NuPage 3-8% w/v Tris-Acetate or 4-12% w/v Bis-Tris gels (Invitrogen) and transferred to polyvinylidene difluoride (PVDF) membranes. The following antibodies were used: rabbit polyclonal antisera against CHK1 (2345, Cell Signaling), CHK1 pS345 (2345, Cell Signaling), cyclin A (SC751, Santa Cruz), CDC2 pY15 (4539,Cell Signaling), FANCD2 (NB100-182; Novus Biologicals), FANCI (A301-254A, Bethyl), FLAG (F7425, Sigma), H3 pS10 (9701, Cell Signaling), pan pS/T-CDK (9477, Cell Signaling), and V5 (13202, Cell Signaling), and mouse monoclonal antisera against α-tubulin (MS-581-PO, Neomarkers).

### Plasmids

The FANCD2 S592A and S592D cDNAs were generated by site-directed mutagenesis of the wild type FANCD2 cDNA using the Quikchange Site-directed Mutagenesis Kit (Stratagene). The forward (FP) and reverse (RP) oligonucleotide sequences used are as follows: S592A FP 5′-CGGCAGACAGAAGTGAAGCACCTAGTTTGACCCAAG-3′; S592A RP 5′-CTTGGGTCAAACTAGGTGCTTCACTTCTGTCTGCCG-3’; S592D FP 5’-GGCGGCAGACAGAAGTGAAGATCCTAGTTTGACCCAAGAG-3’; and S592D RP 5’-CTCTTGGGTCAAACTAGGATCTTCACTTCTGTCTGCCGCC-3’. The full length FANCD2 cDNA sequences were TOPO cloned into the pENTR/D-TOPO (Invitrogen) entry vector, and subsequently recombined into the pLenti6.2/V5-DEST (Invitrogen) destination vector and used to generate lentivirus for the generation of stable cell lines.

### Immunofluoresence microscopy

Hela or Hela FANCD2^-/-^ generated by CRISPR-Cas9 were seeded in at a density of 2 x 10^4^ in chamber slides (Millicell EZ Slide, Millipore) overnight. Cells were treated with 100 nM MMC for 24 h and fixed in fixing buffer (4% w/v paraformaldehyde, 2% w/v sucrose in PBS, pH 7.4) at 4°C for 10 min. Cells were permeabilized using 0.3% v/v Triton-X in PBS for 10 min at room temperature. Cells were blocked in antibody dilution buffer (ADB) (5% v/v goat serum, 0.1% v/v NP-40 in PBS, pH 7.4) and then incubated with mouse-anti H3 pS10 (9706, Cell Signaling) and rabbit-anti-FANCD2. Cells were washed with PBS and incubated in secondary goat-anti-mouse Alexa Fluor 594 and donkey-anti-rabbit Alexa Fluor 488 in ABD for 1 h. Cells were washed in PBS and stained with 4′,6-diamidino-2-phenylindole dihydrochloride (DAPI, Vector Laboratories).

### Cell-cycle synchronization

For early S-phase arrest, cells were synchronized by the double-thymidine block method. Cells were seeded in 10 cm^2^ dishes and treated with 2 mM thymidine (226740050, Acros Organics) for 18 h. Cells were washed with PBS and released into thymidine-free media for 10 h following by a second incubation in 2 mM thymidine for 18 h. Cells were washed twice with phosphate-buffered saline (PBS) and released into thymidine-free media. For M-phase arrest, cells were synchronized using the mitotic shake-off method. HeLa cells were seeded in 10 cm^2^ dishes. Cells were treated with 100 ng/mL of nocodazole (SML1665, Sigma) at 80-90% confluency for 15 h. Mitotic cells were collected by the shake off method and re-plated in nocodazole-free media for cell cycle progression analysis. For late G2-phase arrest, cells were synchronized by reversible inhibition of CDK1 (Vassilev 2006). Cells were seeded in 10 cm^2^ dishes and treated with 6 μM R03306 (15149, Cayman Chemicals) for 16 h. Cells were washed with PBS and released into fresh media without R03306.

### Cell cycle analysis by FACS

Cells were fixed in ice cold methanol, washed in PBS and incubated in 50 μg/mL propidium iodide (PI) (Sigma) and 30 U/mL RNase A for 10 min at 37°C, followed by analysis using a BD FACSVerse flow cytometer. The percentages of cells in G1, S, and G2/M were determined by analyzing PI histograms with FlowJo V10.2 software.

### In-vitro CDK phosphorylation assay

Purified FANCD2 CDK2-Cyclin A proteins were a generous gift from Andrew Deans at the University of Melbourne. In order to remove any previous phosphorylation 2 μg of protein were incubated with 100 U of lambda phosphatase, 10 mM protein metallophosphatases (PMP) and 10 mM MnCl2 at 30°C for 30 min. For the phosphorylation reaction, each reaction tube was composed of the following reagents in this specific order: 10x CDK kinase buffer (250 mM Tris pH 7.5, 250 mM glycerophosphate, 50 mM EGTA, 100 mM MgCl2), 2 μg of protein, 15 nM CDK2:Cyclin A, and 20 μM ATP. Samples were incubated at 30°C for 30 min and the reaction was stopped by adding 10 μL of 10% β-mercaptoethanol in 4x LDS buffer.

### Cell proliferation assay

For cell proliferation assays we performed electrical impedance analysis using the xCELLigence RTCA DP system from Acea Biosciences. Cells were seeded in polyethylene terephthalate (PET) E-plates (300600890, Acea Biosciences). Cells were incubated in the absence or presence of 0.4 μM aphidicolin (APH) or 20 nM mitomycin C (MMC) for 120 h. Electrical impedance measurements were taken every 15 min over the course of the incubation. Statistical analysis was performed using R.

## Acknowledgements

We thank members of the Howlett, Camberg, and Dutta laboratories for critical discussions. We thank Andrew A. Deans and Winnie Tan at the University of Melbourne for purified proteins and Martin A. Cohn at the University of Oxford for HeLa FANCD2^-/-^ cells. This work was supported by NIH/NIGMS grant R01HL149907 (N.G.H.), an American Society of Hematology Bridge Grant (N.G.H.); Rhode Island IDeA Network of Biomedical Research Excellence (RI-INBRE) grant P20GM103430 from the National Institute of General Medical Sciences; and Rhode Island Experimental Program to Stimulate Competitive Research (RI-EPSCoR) grant #1004057 from the National Science Foundation. We declare that we have no conflicts of interest.

---

**Figure S1.**
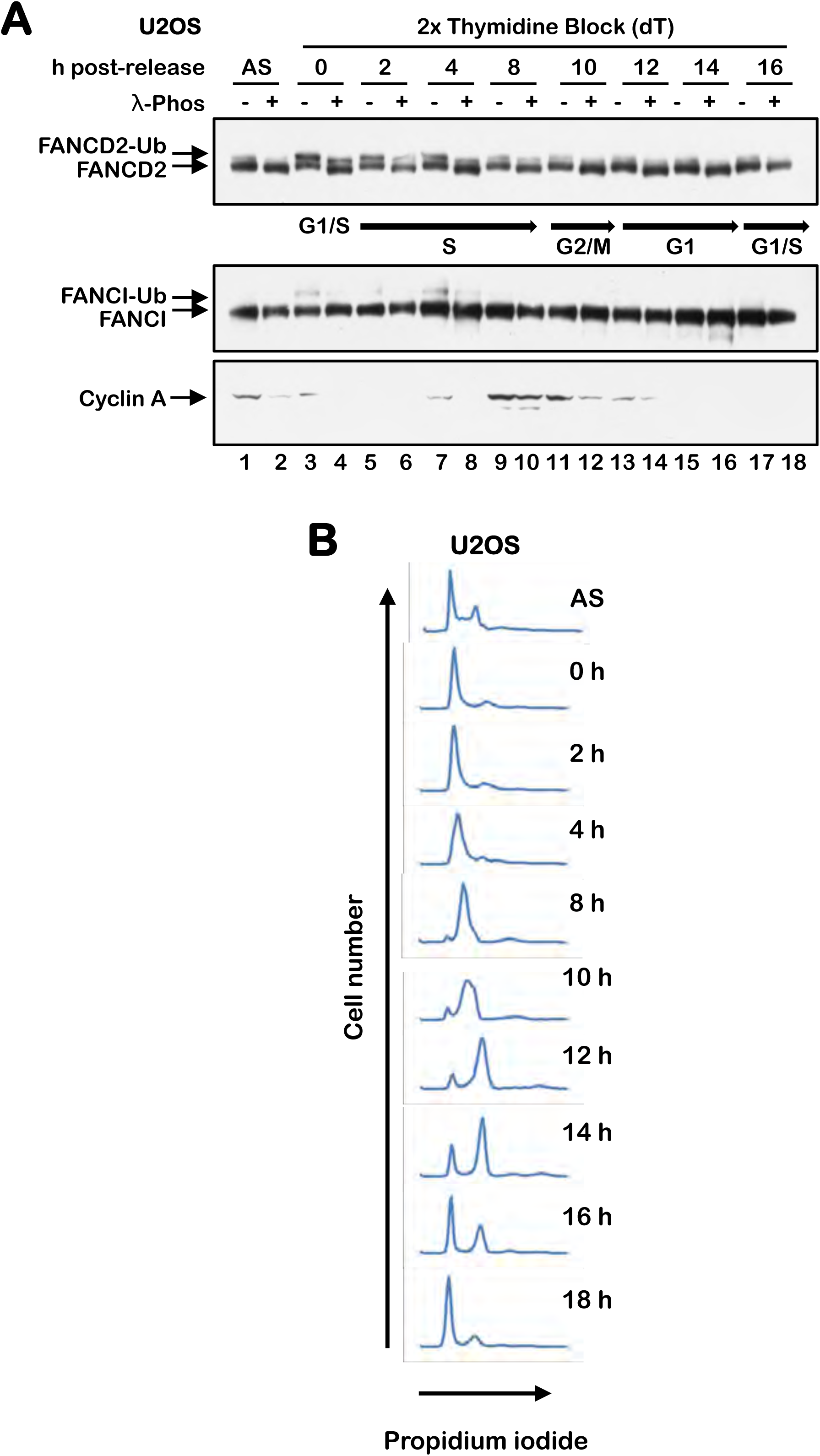
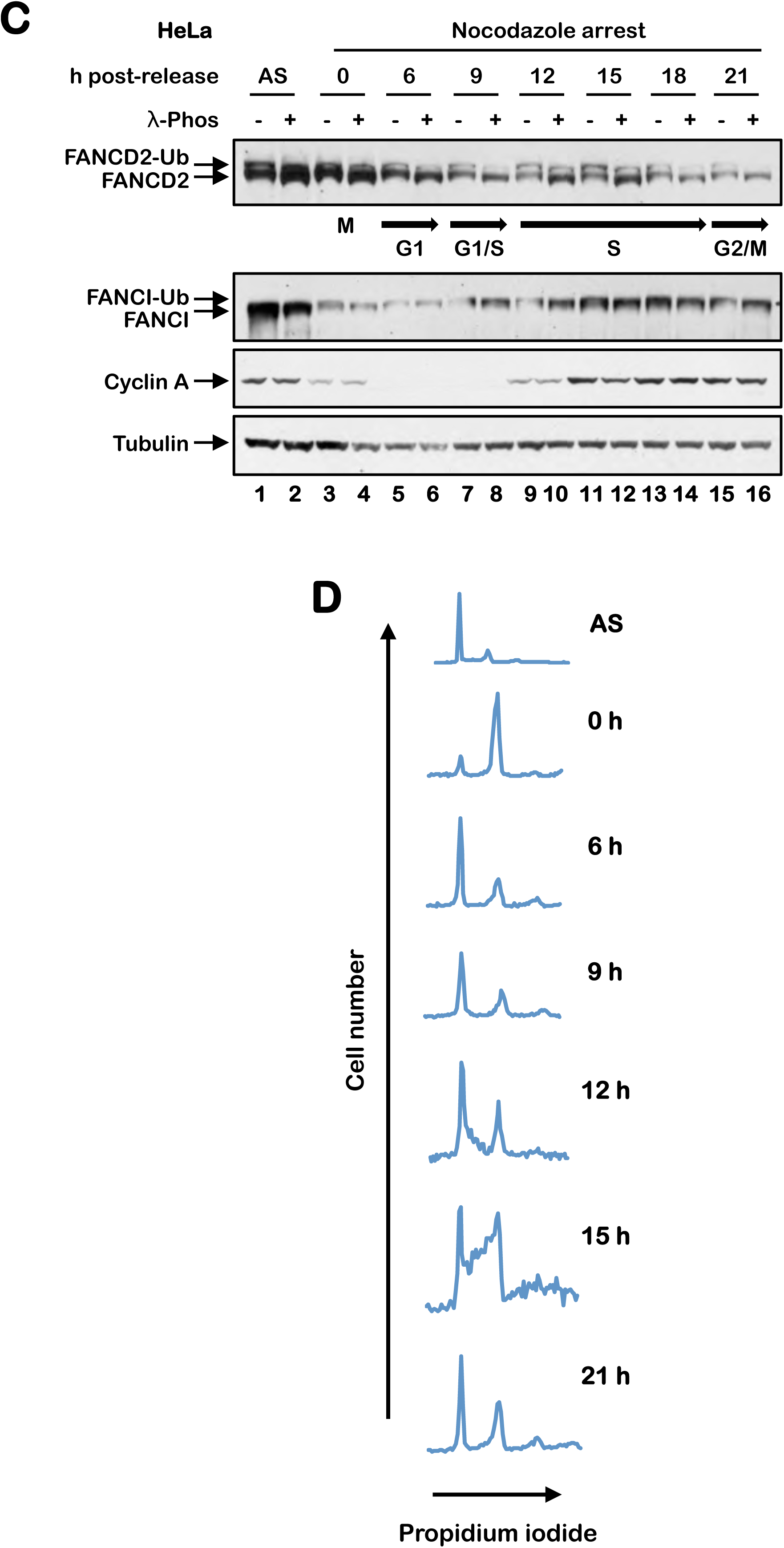
Phosphorylation of FANCD2 during S-phase of the cell cycle. (A) U2OS cells were synchronized at the G1/S boundary by a double-thymidine block and released into thymidine-free media. Cells were harvested, lysed in lambda phosphatase buffer and incubated in the presence or absence of lambda phosphatase. Samples were analyzed by immunoblotting. (B) Cells were fixed, stained with propidium iodide and analyzed by flow cytometry to determine cell cycle stage. (C) HeLa cells were synchronized in M-phase with a nocodazole block. Mitotic cells were physically detached by the shake-off method and released into nocodazole-free media. Cells were harvested, lysed in lambda phosphatase buffer and incubated in the presence or absence of lambda phosphatase. Samples were analyzed by immunoblotting. (D) Cells were fixed, stained with propidium iodide and analyzed by flow cytometry to determine cell cycle stage.

**Figure S2.**
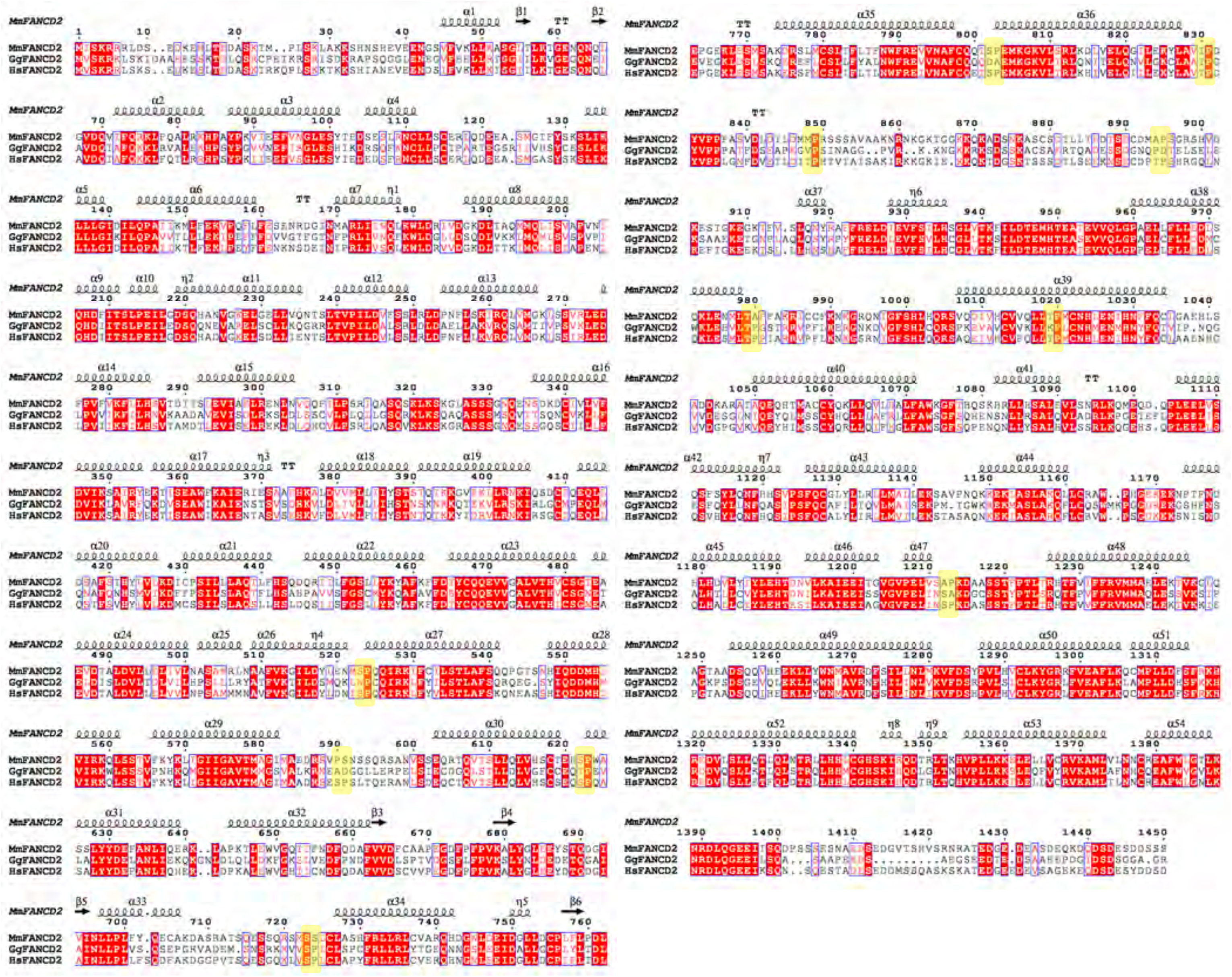
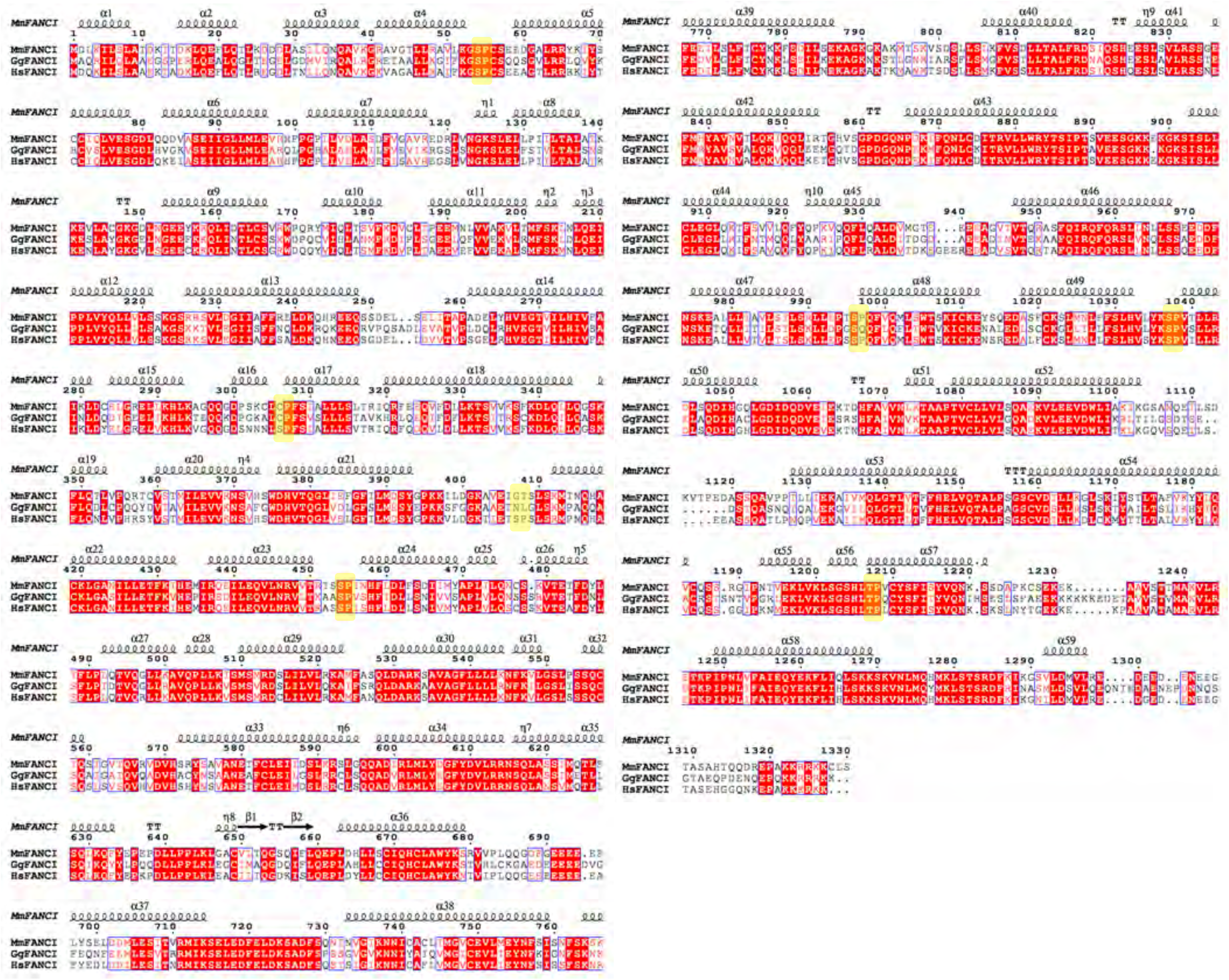
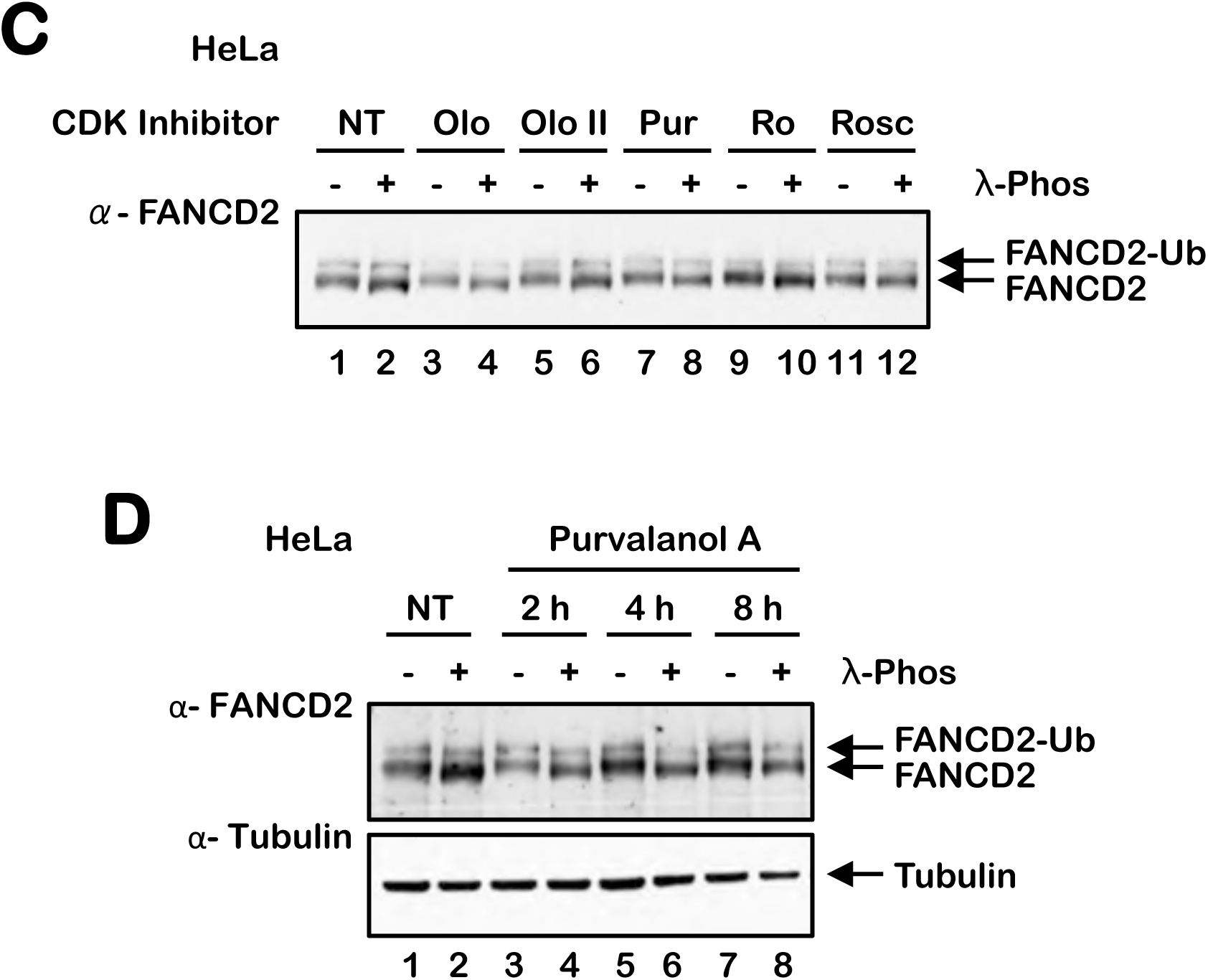
FANCD2 is a CDK substrate. (A-B) Shown is an alignment of mouse, human, and chicken FANCD2 (A) and FANCI (B) amino acid sequences using the T-Coffee server, with the secondary structure of mouse FANCD2 or FANCI illustrated. Putative CDK phosphorylation sites are shaded in yellow. (C) HeLa cells were incubated in the absence (NT) or presence of 10 μM olumucine (Olo), 10 μM olomucine II (Olo II), 10 μM purvalanol A (Pur), 10 μM RO3306 (Ro) and 20 μM roscovitine (Rosc) for 24 h. Whole cell lysates were incubated in the absence or presence of lambda phosphatase and analyzed by immunoblotting. (D) HeLa cells were treated with 10 μM purvalanol A for the indicated times, lysates were incubated in the absence or presence of lambda phosphatase, and analyzed by immunoblotting.

**Figure S3.**
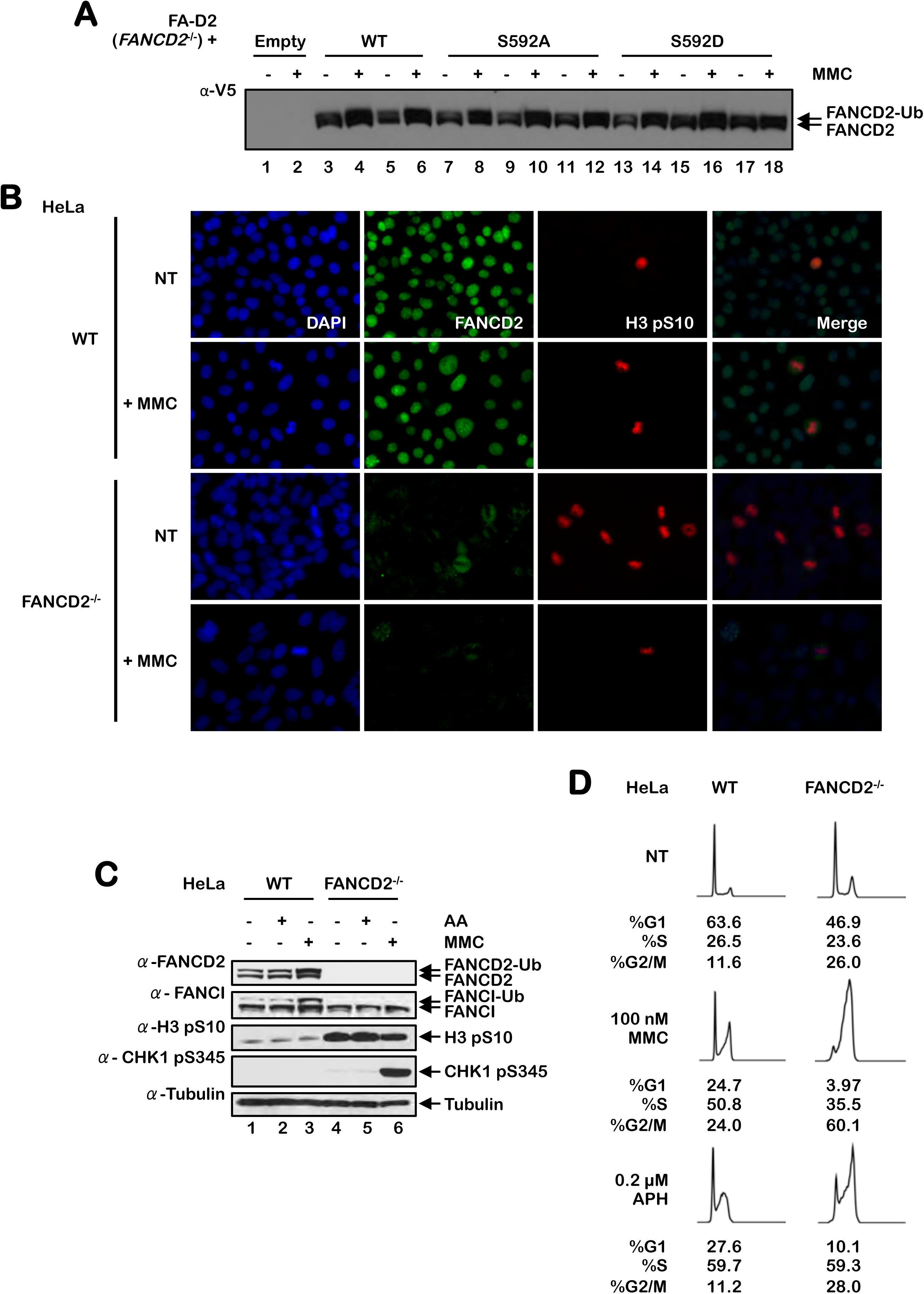
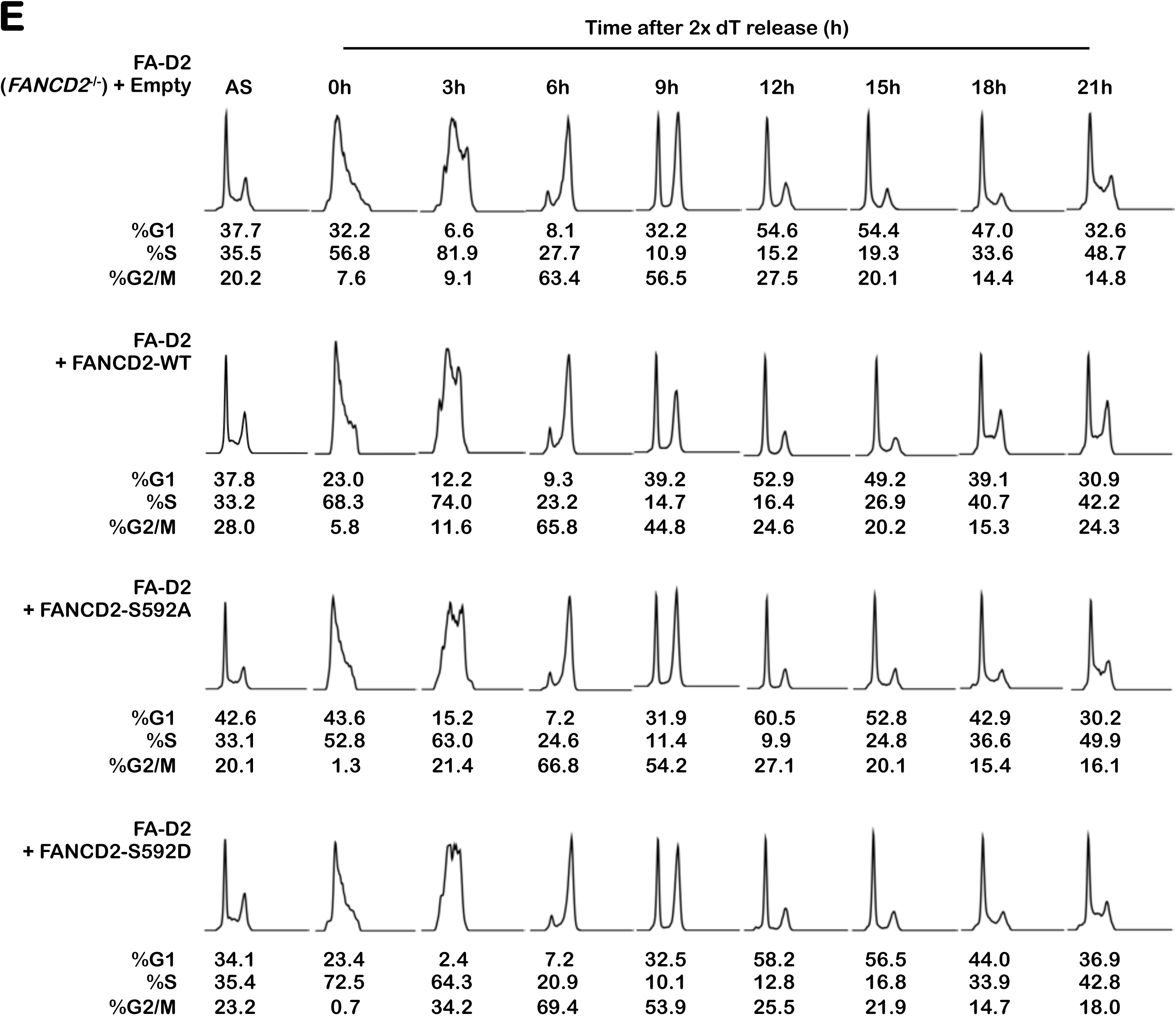
Mutation on S592 affects cell cycle progression / Cells lacking FANCD2 exhibit increased mitotic arrest in unstressed conditions. (A) Polyclonal populations of FA-D2 (FANCD2^-/-^) patient cells expressing empty vector or V5-tagged FANCD2-WT, FANCD2-S592A, or FANCD2-S592D were incubated in the absence (-) or presence (+) of 200 nM mitomycin C (MMC) for 24 h and whole-cell lysates were analyzed by immunoblotting. (B) Wild-type HeLa and HeLa FANCD2^-/-^ cells generated by CRISPR-Cas9 gene editing were incubated in the absence or presence of 100 nM MMC for 24 h. Cells were fixed and co-immunofluoresence microscopy was performed for FANCD2 (green) and H3 pS10 (red). (C) The same cells were treated with 100 nM MMC or 1 mM acetaldehyde for 24 h and whole-cell lysates analyzed by immunoblotting. (D) Cells were incubated in the absence or presence of 100 nM MMC or 0.2 μM APH for 24 h. Cells were fixed, stained with propidium iodide and analyzed by flow cytometry. Cell cycle stage analysis was performed using the FlowJo V10.2 software. (E) FA-D2 cells stably expressing empty vector or V5-tagged FANCD2-WT, FANCD2-S592A, or FANCD2-S592D were synchronized at the G1/S boundary by double-thymidine block (2x dT), and then released into thymidine-free medium. At the indicated time points, cells were fixed, stained with propidium iodide and analyzed by flow cytometry. Cell cycle stage analysis was performed using the FlowJo V10.2 software.

**Figure S5.**
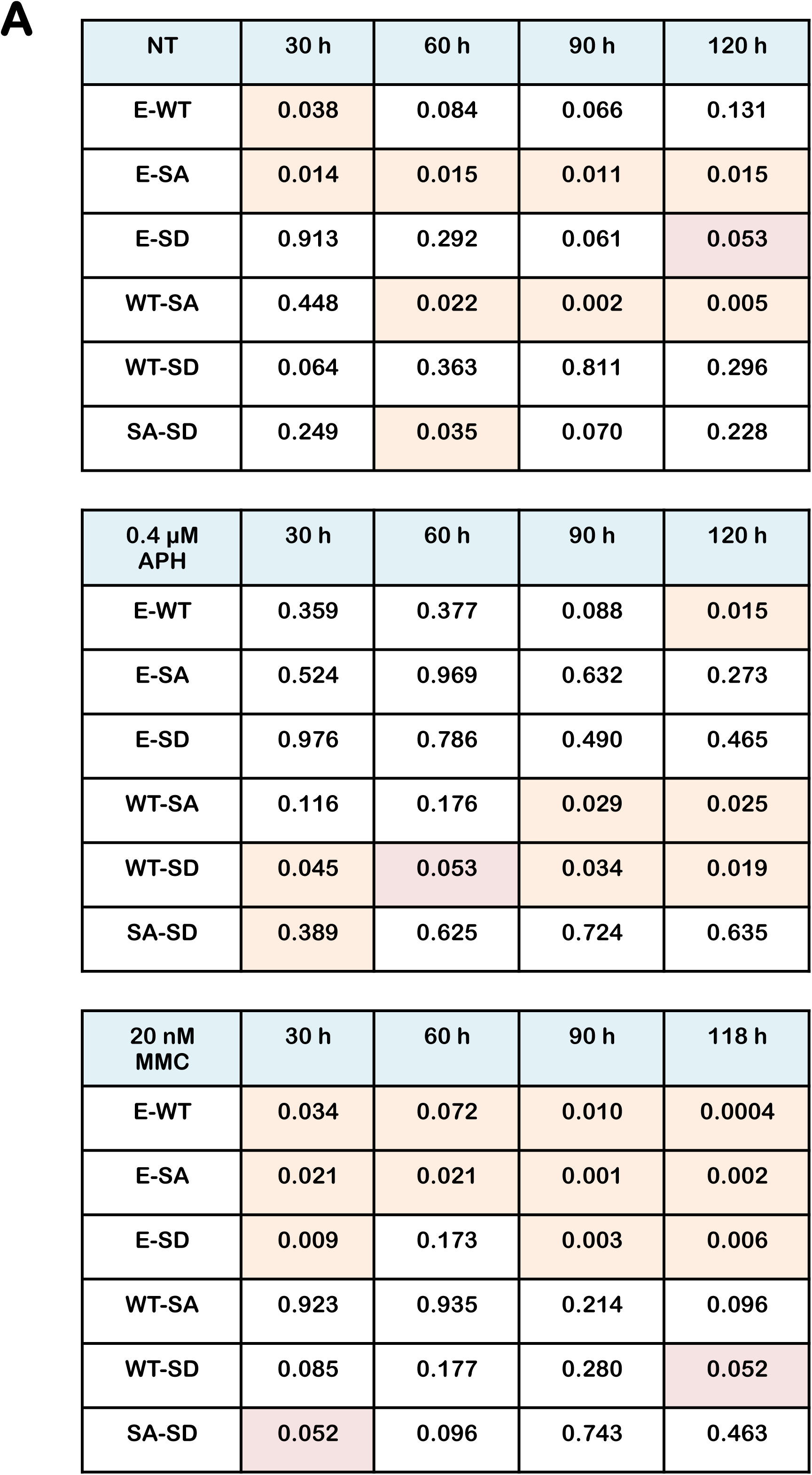
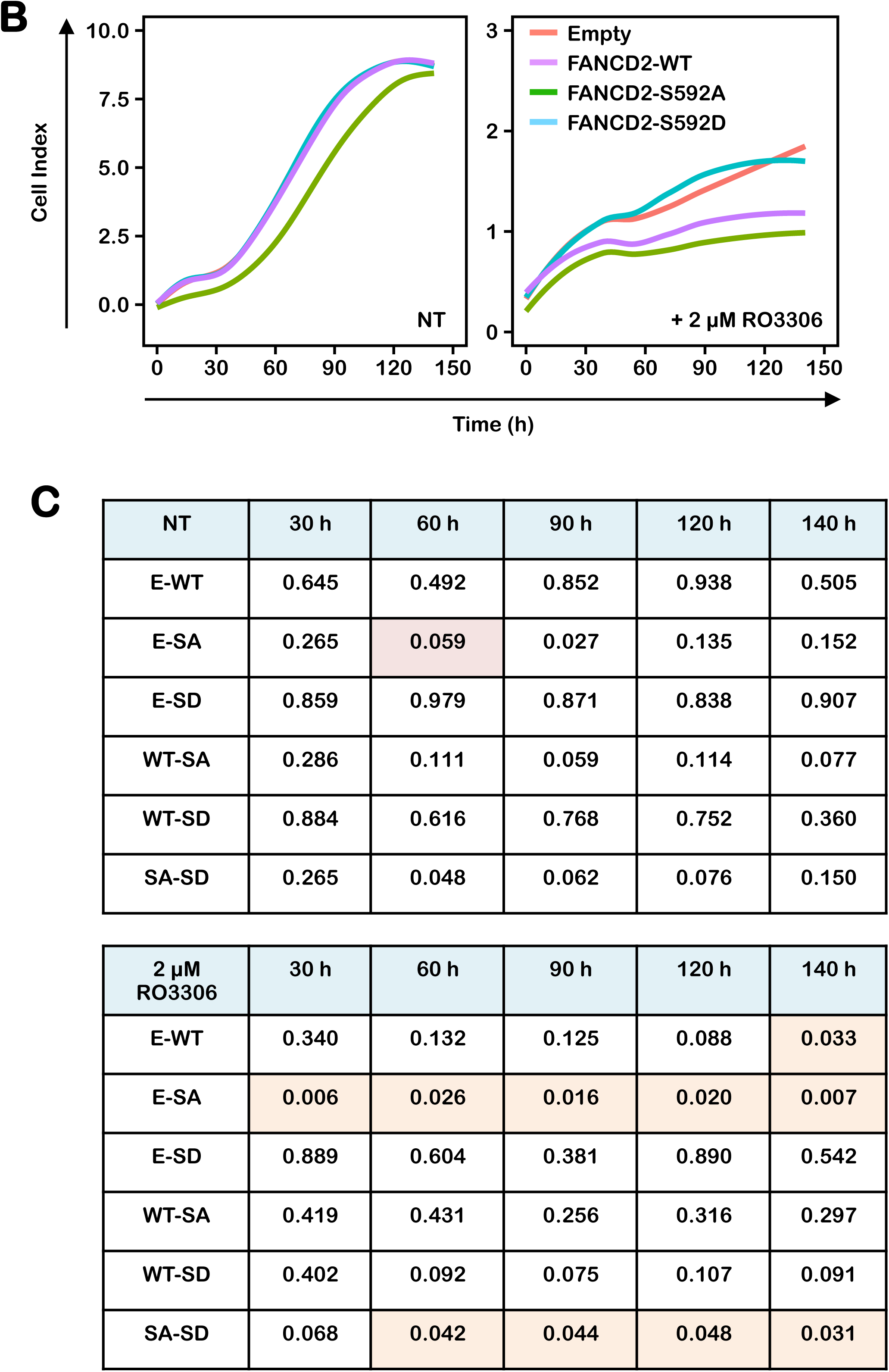
Mutation of FANCD2 S592 leads to decreased proliferation under conditions of replicative stress. (A) FA-D2 cells stably expressing empty vector or V5-tagged FANCD2-WT, FANCD2-S592A, or FANCD2-S592D were incubated in the absence (NT) or presence of 0.4 μM aphidicolin (+APH) or 20 nM mitomycin C (+MMC). Cellular proliferation was monitored by measuring electrical impedance every 15 minutes over a 120 h period using the xCELLigence real time cell analysis system. Student’s t-test was used to compare electrical impedance measurements between populations at 30 h, 60 h, 90 h and 120 h. (B) The same cells were incubated in the absence or presence of 2 μM RO3306 and cellular proliferation was monitored by measuring electrical impedance every 15 minutes over a 140 h period using the xCELLigence real time cell analysis system. (C) Student’s t-test was used to compare electrical impedance measurements between populations at 30 h, 60 h, 90 h, 120 h, and 140 h.

